## Supplementary material for "BARcode DEmixing through Non-negative Spatial Regression (BarDensr)": S1 Appendix

[1]Mortimer B. Zuckerman Mind Brain Behavior Institute, Columbia University, New York, New York, United States of America
[2]Department of Statistics, Columbia University, New York, New York, United States of America
[3]Center for Theoretical Neuroscience, Columbia University, New York, New York, United States of America
[4]Grossman Center for the Statistics of Mind, Columbia University, New York, New York, United States of America
[5]Department of Neuroscience, Columbia University, New York, New York, United States of America
[6]Department of Systems Biology, Columbia University, New York, New York, United States of America
[7]Data Science Institute, Columbia University, New York, New York, United States of America
[8]Cold Spring Harbor Laboratory, Cold Spring Harbor, New York, United States of America
[*]

### A Data preprocessing

The data is preprocessed before input into the model as follows: first the data was max-projected across all z-stacks. The channel color-mixing was corrected and the background was removed using rolling-ball background subtraction [1]. Then the different image stacks were registered to the same voxels, using the Image Alignment Toolbox (ECC image alignment algorithm) [2].

Next, we performed a crude noise-normalization on each frame. First we estimated the noise level on each frame by spatially high-pass filtering (i.e., original image minus a Gaussian-filtered image, with a sigma of 2 voxels) to isolate spatially-uncorrelated noise, then computing the standard deviation. (See e.g. [3] for a related approach applied in the temporal domain.) Then we divided each original frame by its estimated noise scale to obtain the noise-normalized images.

When necessary, we used an additional step to normalize the image intensities. We found this step reduced the variations across the spots and image frames. Specifically, a Gaussian filter with a sigma of 5 voxels (which is approximately twice as big as the typical rolony size) was applied to each image frame, yielding a blurred image. The normalized image was obtained by dividing each original image frame with this blurred image.

### B Supplementary video

The supplementary videos at https://tinyurl.com/y7zzyrd4 correspond to S7-S9 Figs, with the image colors indicating the channel intensity. The first panel is the original data; the second panel is the reconstruction from BarDensr; the third panel is the reconstruction from BarDensr after sparsifying the low-intensity voxels (those smaller than the maximum intensity of the unused barcodes are forced to be zero) and then updating the non-zero values in $\mathbf{F}$ based on the other re-fit parameters; the fourth and fifth panels are the residual of the reconstructions from the second and third panels, respectively. The color scales are the same across the first three panels, and also across the last two panels.

### C NeuroCAAS input file format

NeuroCAAS inputs include the following: the stack of images in HDF5 format; the configuration file in YAML format; the codebook in CSV format.

For the HDF5 file, each image name should be formatted as 'round_r, channel_c', where 'r' and 'c' start from 1 and are the indices of round and channel for each image. Each image should be formatted as a 3D image (e.g., $(2048 \times 2048 \times 1)$ for a 2D image with 2048 voxels in X and Y directions), and they must be preprocessed and properly registered across rounds and channels before input to this platform.

The YAML file is used to specify the parameters used for the analysis, as well as the information on the codebook corresponding to the input data (see below). The details for each argument are as follows:

- 'blur_level': indicates the expected size of the rolonies, passed to the point-spread function $\mathbf{K}$ in Table 1.
- 'downsample_level': scale of the downsampling for coarsening process. The three integers indicate that for each coordinate, how many voxels should correspond to a single voxel in the original image, after the downsampling.
- 'tile_size': the size of the tiles for patching. Each tile will be analyzed independently in the following 'fine' process, and the barcodes that are absent in each tile will be removed.
- 'detection_thre': between 0 to 1, indicating what percentage of the maximum intensity of unused barcodes is used to determine the threshold. This controls the threshold for removing the absent barcodes in each tile, as well as the threshold for detecting the blobs.
- 'lam': sparsity penalty corresponding to $\lambda$ in Section L. This value is highly dependent on the data and the preprocessing steps. For a noise-normalized images, we used $\lambda = 0.2$.
- 'codebook_name': specify the name of the codebook file uploaded (see below).
- 'unused_included' and 'num_unused': these two arguments specify if and how many of the unused barcodes are included in the uploaded codebook. Unused barcodes are necessary for the analysis to determine the threshold to detect the absent barcodes in each tile, as well as to detect blobs. Note if 'unused_included' is *yes* and 'num_unused' is 3, BarDensr assumes that the unused barcodes are the

last three barcodes in the codebook. If 'unused_included' is *no* and 'num_unused' is 2, BarDensr will generate two unused barcodes for the analysis.

The CSV file stores the binary barcodes (second column) of each gene (first column). The binary barcodes should be channel-major order (i.e., r1_c1, r1_c2, ..., r2_c1, r2_c2.., ).

### D Hardware time and cost comparisons

To develop an efficient implementation of BarDensr on the NeuroCAAS cloud platform [4], we needed to find the most cost-effective hardware for the job. Using a $1000 \times 1000$ sized image from the experimental data described in the main text, we ran the model on several different AWS instance types. The most cost-effective machine was `m5.2xlarge`, which completed the analysis in three and a half minutes with a total cost of two cents. On the other extreme, the `p3.2xlarge` machine completed the analysis in one minute with a total cost of five cents. As a compromise between speed and cost, we settled on the `p2.xlarge` machine, which completes the analysis in two minutes with a total cost of three cents.

### E Single Round Matching (SRM) and 'hand-curated' method

We compared BarDensr against several different alternative methods, including one we call 'SRM.' This method is an implementation of the widely-used 'blobs-first' algorithms suggested in the literature [5, 6]. First, blobs were detected in every channel in the first round by finding local maxima on a per-channel basis. These blobs were then used as a reference in understanding subsequent rounds (the first round is used as the reference since it usually has the least corruption by noise and artifacts, such as phasing and photo-bleaching).

At each detected rolony position, SRM then read out the signal intensities from all channels/rounds as a vector of length $R \times C$. This vector was compared against each barcode in the library. Each detected rolony was assigned to the barcode with the greatest similarity (as measured by a dot-product). For some rolonies the similarity was low to all barcodes; these rolonies were filtered out. Thresholds were determined using the Bayesian optimization method from [7].

The 'hand-curated' method in the main text corresponds to SRM described here. After the process of SRM with the chosen threshold, we manually checked the detected rolonies and the assigned barcodes to make sure that the results were reasonable.

### F Correlation-based method

We also compared BarDensr against a 'correlation-based method' [8, 9]. This approach begins by computing a vector of length $R \times C$ for every voxel, indicating the fluorescence signal in each round and channel at that voxel. At each voxel, for each barcode, the cosine distance between this vector and the barcode was computed. The barcode with the minimum cosine distance was assigned to be a *potential* gene identity for each voxel. Finally, a 'minimum distance image' was constructed: for each voxel, this image contains the cosine distance between that voxel's $R \times C$ vector and the barcode which it is most similar to. Coordinates of blobs were found by a seeking local minima in this image. Thresholds were again determined via [7].

For the experimental data used in Fig 4D, we added additional preprocessing steps in order to remove the unwanted noise. Specifically, signal intensity at each voxel location $m$ for frame $r, c$ is computed as $\frac{\mathbf{X}_{m,r,c}}{P + \left(\sum_r^R \mathbf{X}_{m,r,c}^2\right)^{\frac{1}{2}}}$, before computing the dot products. $P$ is a constant that is data-dependent, and represents the magnitude of the noise floor of the data.

### G Simulation

#### Generating arbitrary distribution for genes

For the simulation benchmarking in Fig 3A and S4 Fig, we used a set of barcodes that is similar to the ones used in STARmap experiment [5], with total of 57 genes. This simulated data is similar to our original experimental data in that it has six rounds and four channels in total, and the scale of the number of barcodes is also similar. This simulated data was chosen instead of our experimental data in order to directly apply *starfish* method (the *starfish* application on our original experimental data was not available at the time this analysis was conducted). In creating simulations, we wanted to accurately represent the uneven distribution of genes; in real data some genes are more abundant than others. Therefore, we began by randomly selecting 10 out of 57 genes to be 'abundant' genes. In generating a dataset with simulated rolonies, we created rolonies with these abundant genes roughly ten times more often than the other rolonies.

#### Dropout

We used two setups to generate simulated testing data: without dropout and with dropout. In the experimental data, it is commonly observed that a small portion of rolonies disappear/diminish in some rounds. (Based on our visual inspection of the experimental data, qualitative dropout events were observed in $< 5\%$ of the rolonies detected in this data, but we did not attempt to estimate this dropout rate precisely.) In the 'Dropout' simulations, we tried to mimic this phenomenon.

Specifically, for the 'no dropout' simulations, we generated the data with the following process. 1. Generate the spot position with a uniform distribution across the voxels. 2. For each spot position, generate the spot identity (gene) using a prespecified gene distribution (as discussed above). 3. For each position $m$ and gene $j$ pair from steps 1 and 2, the *magnitude* of the rolony density at $(m, j)$ was generated from a uniform distribution in the range $(10, 40)$. We use these values to fill in the rolony density, $\mathbf{F}$. 4. We then generate synthetic data according to the BarDensr model. Finally, we add some speckle noise. Note that in our simulation, parameters such as the per-frame intensity ($\alpha$), phasing ($\rho$), and color-mixing ($\varphi$) were left out for simplicity.

For the 'dropout' simulations, 50% of the simulated spots were randomly selected to be the 'dropout spots.' For each 'dropout spot', one round is selected randomly and the signal intensity for that spot for that round is diminished to 10% of the original signal. The simulation process is otherwise the same.

#### Hybrid simulation

To test the efficacy of our model on even more realistic data, we used a 'hybrid simulation,' as in e.g. [10]. In essence, the hybrid simulation creates a fake dataset by superimposing the real data with additional synthetic rolonies to the data. The question is whether the algorithm can at least find the synthetic rolonies which were added. The results are shown in Fig 3B. In generating the synthetic rolonies, we used the codebook used in the original experiment, and the genes of the synthetic rolonies were given by the observed gene distribution from the hand-curated analysis of the same dataset.

We ran a few different versions of these simulations. There were several key parameters which we varied:

- We had both 'dropout' and 'no dropout' versions; in the dropout versions some of the synthetic rolonies had signal in one round diminished.
- We could vary the number of synthetic spots which were injected ($S$). According to the hand-curated analysis of the real data, the real data contained approximately 400 spots in this field of view. We investigated how the number of spots affected the results, looking at $S \in \{30, 100, 500, 2000\}$.
- We could vary the intensity of the synthetic spots, relative to the maximum intensity observed in the data. We varied this between 10% and 90%.

### H Error analysis

In simulated data, we can exactly quantify the different kinds of errors that BarDensr makes, by comparing against the true rolony positions used to make the simulated data. We first examined the 'total hit rate' in our Receiver Operating Characteristic curve (ROC curve, shown as the dotted lines in Fig 3A). For this ROC, we consider a spot to be successfully detected by the algorithm as long as the algorithm finds *any rolony* near the site of a true rolony – even if the algorithm incorrectly assigns the gene associated with that true rolony. The ROC for BarDensr clings closely to the upper left side of the plot, suggesting nearly perfect performance. We also looked at what we call the 'hit rate' – for this ROC we consider a spot to be successfully detected only if the algorithm detects a rolony in the right place and of the right gene. Fig 3A shows the results, suggesting most errors were caused by gene mis-identification.

### I Cell segmentation

For Fig 4C, we first segmented the cells in the selected region with the following process. We first obtained the max projection across $R \times C$ frames from the image stacks. After applying a Gaussian filter with a sigma of 8 voxels, all the voxels with intensity lower than 10% of the maximum intensity were assigned to be zero. Finally, we used a watershed segmentation algorithm to identify contiguous cellular regions. This results in 47 segmented cells in the region. Four of these occupied less than 100 voxels in total and were removed from the analysis. The cell segmentation results for Fig 6 were obtained from the previous study [11].

### J Benchmark experimental data

For the benchmark data used in Fig 4D, we selected a region from mouse motor cortex section from a recent study [11] as explained in the main text (Methods section). This region in the cortex consists of 35 fields of view. We manually inspected one field of view, and selected a $200 \times 200$ region that is relatively dense. In order to find every last rolony in this region, we used several approaches. We used a hand-curated approach, a correlation-based method, and BarDensr on this region. Wherever there was any disagreement we determined the truth by direct inspection. We found 134 rolonies in total.

The methods used to obtain Fig 4D are similar to what have been described in Sections E and F. For *starfish*, we used both 'spot-based' and 'pixel-based' approaches. The latter is similar to the correlation-based method described in Section F. To plot the ROC for BarDensr and the correlation-based method, we only needed to change the threshold for the blob detection. For each of the two approaches of *starfish*, we have changed two parameters and kept all other parameters fixed as the default values.

### K Sparsifying and coarse-to-fine

#### K.1 Handling tens of thousands of barcodes with sparsifying and coarse-to-fine

To scale up BarDensr, we tested if eliminating unnecessary barcodes could help accelerate computation. For this purpose, we set up a simulation with 53,000 barcodes and 17 sequencing rounds, similar to the setup in a larger scale experiment such as [12] (S6 Fig). We generated a dataset with $50 \times 80$ voxels and a total of 40 spots. We then processed this data in two steps.

In the first step (the 'coarse' step), the image was five times downsampled, and BarDensr was applied to the downsampled data. For each gene, if the maximum intensity for a rolony density was lower than $10^{-5}$, that gene was considered to be absent.

In the second step (the 'sparsified fine' step), we then applied BarDensr to the original, full-resolution data – but only using those barcodes that weren't 'absent' in the previous step. To make this approach even faster, we also used the learned parameters from the downsampled data as initial conditions for the algorithm's run on the full-resolution data. Moreover, in this second step, the parameter $b$ and $\alpha$ were not updated at all, since we found that they were learned quite accurately in the first step.

After the first step, 71 out of 53,000 barcodes were kept to be used in the second step. This sparsified coarse-to-fine approach sped up our analysis by more than a factor of 10.

##### K.2 Coarse-to-fine

The method described above involves both sparsifying and using a kind of coarse-to-fine approach. We also investigated performance using only the coarse-to-fine aspect. These investigations are summarized in Fig 5A. First, a $1000 \times 1000$ image was simulated with $20,000$ spots following the simulation process described above, with no dropout (Section G). The image was then downsampled to $500 \times 500$ and BarDensr was applied to this downsampled data to estimate $\mathbf{F}, \alpha, a$ and $b$, with 20 iterations. These parameters were then used as the initial conditions to run the model with the full size image (note that in order to use the parameters from the downsampled image to initialize the full-resolution model, we needed to upsample $\mathbf{F}$ and $a$). Finally, we tried running the model directly on the full-resolution data (without using the downsampled data to get initial conditions). We then compared the results. Both approaches work better if they are allowed to run longer, because they use an iterative approach to optimize the loss function. Eventually, both approaches yield the same results. However, Fig 5A shows that the coarse-to-fine approach is able to achieve the best possible performance three times faster.

##### K.3 Sparsifying leads to speedups even in the case of a small barcode library

We also tested the sparsifying approach on the experimental data (Fig 5B). The original $1000 \times 1000$ image was first five times downsampled to obtain a $200 \times 200$ 'coarse' image, and the parameters were learned from this downsampled image ('coarse' process). After five times upsampling of learned $\mathbf{F}$ to obtain the image on the original scale, both the original image and the upsampled $\mathbf{F}$ were split into $4 \times 4$ patches. Each patch is of size $250 \times 250$ (plus 20 voxel edges in the end of both dimensions, whenever the coverage of the patch does not exceed the image region). These 16 patches cover the entire $1000 \times 1000$ image with overlaps on the edge regions. For each patch, the barcodes that have the maximum intensity lower than the maximum intensity of the two unused barcodes in the upsampled $\mathbf{F}$ were considered to be absent from the region. Each patch was then used to fit the model at the original scale to learn $\mathbf{F}$ and $a$, but using a smaller amount of barcodes for the binary codebook matrix $\mathbf{B}$. For this 'fine' process, the parameter $b$ and $\alpha$ were not updated but the ones learned from the coarse process were used. After the model was run on all the 16 patches, we needed to stitch the results back together into a single result for the entire field of view. This is slightly involved, because the patches concerned overlapping regions of voxels. Indeed, we ensured that the border-regions between any two patches contained 20 voxels of overlap. For each voxel in these overlap regions, we used the signal from the patch whose center was closest to that voxel. The results were compared to the 'fine only' approach. We filled $\mathbf{F}$ with zero for the removed barcodes in each patch, in order to keep the original dimension for the following process.

In order to compare the agreement between 'sparsifying' approach and the 'fine only' approach, we computed two ROC curves. In each curve, one of the results was used as the gold standard for the other. Specifically, when using the 'fine only' as the gold standard, a threshold was determined based on the maximum intensity of the unused barcodes in the 'fine only' results, and a binary matrix of the size of the original image ($1000 \times 1000$) was generated for each barcode ('0' indicates there is no signal, '1' indicates there is signal, for each voxel), which is the final gold standard to compare with the sparsifying result. The same process applies when using the sparsifying result as the gold standard. For computing ROC, rolonies within 3 voxel radius of $\mathbf{F}$ with the same barcode $j$ were considered to belong to the same rolony.

#### L Algorithm Details

The primary computational challenge of this method is to solve a constrained optimization problem.

$$\begin{aligned} \min_{\theta \in \Theta} \quad & L_{\text{sparsity}}(\theta) \\ \text{subject to} \quad & L_{\text{reconstruction}}(\theta) \leq \omega, \end{aligned}$$

where

$$
\begin{aligned}
L_{\text{reconstruction}}(\theta) &\triangleq \sum_{m,r,c}\left(\mathbf{X}_{m,r,c}-\left(a_m+b_{r,c}+\alpha_{r,c}\sum_{j,m',c'}^{J,M,C}\mathbf{K}_{m,m'}\mathbf{F}_{m',j}\varphi_{c,c'}\mathbf{Z}_{r,c',j}\right)\right)^2\\
L_{\text{sparsity}}(\theta) &\triangleq \sum_{m,r,c,c',j}\alpha_{r,c}\mathbf{F}_{m,j}\varphi_{c,c'}\mathbf{Z}_{r,c',j}\\
\theta &= (\mathbf{F},\rho,\alpha,b,a,\varphi)\,.
\end{aligned}
$$

Here $\Theta$ denotes the set of feasible parameters; in our case, $\Theta$ simply requires that all variables are at least $10^{-10}$. The threshold $10^{-10}$ was chosen somewhat arbitrarily and serves to ensure numerical stability of the optimization process. Technically we also believe that $\rho_c < 1$ for every $c$, but in practice we found it unnecessary to enforce this constraint.

We approach the reconstruction constraint using Lagrange multipliers. We define:

$$
\begin{aligned}
\mathcal{L}(\theta,\lambda) &= L_{\text{reconstruction}}(\theta)+\lambda L_{\text{sparsity}}(\theta)\\
\theta^*(\lambda) &= \arg\min_{\theta\in\Theta}\mathcal{L}(\theta,\lambda)\\
L^*_{\text{reconstruction}}(\lambda) &= L(\theta^*(\lambda)).
\end{aligned}
$$

Assuming we can evaluate $L^*_{\text{reconstruction}}(\lambda)$, we can solve the overall constrained optimization problem by taking

$$
\lambda^* = \max\left\{\lambda:\ L^*_{\text{reconstruction}}(\lambda)\le\omega\right\}
$$

and taking our final parameters to be $\theta^*(\lambda^*)$. It is unclear whether strong duality holds in this case, so the resulting parameters may not be optimal. However, in practice we find that they give useful results for uncovering rolonies.

In conclusion, to solve our overall problem it suffices to be able to solve $\min_{\theta\in\Theta}\mathcal{L}(\theta,\lambda)$ for any fixed $\lambda$. We approach this problem via a blockwise coordinate descent approach. Specifically, we start with an initial guess and iterate through a variety of updates until convergence is achieved. Throughout, we will use the notations

$$
\begin{aligned}
[x]_+ &= \begin{cases} x & \text{if } x > 10^{-10}\\ 10^{-10} & \text{otherwise}\end{cases}\\
\texttt{xmab}_{m,r,c} &= \mathbf{X}_{m,r,c}-a_m-b_{r,c}\\
\mathbf{X}^{(j)}_{m,r,c} &= \mathbf{X}_{m,r,c}-a_m-b_{r,c}-\alpha_{r,c}\sum_{j'\neq j}\sum_{m',c'}\mathbf{K}_{m,m'}\mathbf{F}_{m',j'}\varphi_{c,c'}\mathbf{Z}_{r,c',j'}\\
\tilde{\mathbf{F}}_{m,j} &= \sum_{m'}\mathbf{K}_{m,m'}\mathbf{F}_{m',j}.
\end{aligned}
$$

$\alpha$ **update.** For each $r,c$, the relevant portion of the loss for $\alpha_{r,c}$ is given by

$$
\begin{aligned}
\mathcal{L}(\alpha_{r,c}) = \sum_m&\left(\mathbf{X}_{m,r,c}-\left(a_m+b_{r,c}+\alpha_{r,c}\sum_{j,c'}\tilde{\mathbf{F}}_{m,j}\varphi_{c,c'}\mathbf{Z}_{r,c',j}\right)\right)^2\\
&+\lambda\sum_{m,c',j}\alpha_{r,c}\mathbf{F}_{m,j}\varphi_{c,c'}\mathbf{Z}_{r,c',j}+\cdots.
\end{aligned}
$$

Fixing all other variables, subject to the constraint that $\alpha_{r,c}\ge 10^{-10}$, the lowest possible value of this loss is given by

$$
\alpha_{r,c}\leftarrow\left[\frac{\sum_m\left(\texttt{xmab}_{m,r,c}\sum_{j,c'}\tilde{\mathbf{F}}_{m,j}\varphi_{c,c'}\mathbf{Z}_{r,c',j}\right)-\sum_{m,c',j}\mathbf{F}_{m,j}\varphi_{c,c'}\mathbf{Z}_{r,c',j}}{\sum_m\left(\sum_{j,c'}\tilde{\mathbf{F}}_{m,j}\varphi_{c,c'}\mathbf{Z}_{r,c',j}\right)^2}\right]_+.
$$

Note that this update can be done in parallel across all $r,c$.

$\rho$ **update.** We update the variable $\rho$ via a line-search. One at a time, we look at $\rho_c$ and consider possible values for this parameter in the interval $[\rho_c/2, 3\rho_c/2]$. We search for values of $\rho_c$ in this interval which minimizes the loss.

$a, b$ **updates.** Fixing all other variables, the best possible values for $a$ are easy to find. The same goes for $b$. These values are given by

$$a_m \leftarrow \left[ \frac{1}{RC} \sum_{r,c} \left( \mathbf{X}_{m,r,c} - b_{r,c} - \alpha_{r,c} \sum_{j,c'} \tilde{\mathbf{F}}_{m,j} \varphi_{c,c'} \mathbf{Z}_{r,c',j} \right) \right]_+$$

$$b_{rc} \leftarrow \left[ \frac{1}{M} \sum_{m} \left( \mathbf{X}_{m,r,c} - a_m - \alpha_{r,c} \sum_{j,c'} \tilde{\mathbf{F}}_{m,j} \varphi_{c,c'} \mathbf{Z}_{r,c',j} \right) \right]_+ .$$

$\mathbf{F}$ **update.** We make updates to $\mathbf{F}$ one column at a time. We select a random column, $j^*$, and then update the values of $\{\mathbf{F}_{m,j^*}\}_{m \in \{1 \cdots M\}}$. We update these values via a projected coordinate descent algorithm. Let $f$ denote the $j$th column of $\mathbf{F}$ and define

$$\begin{aligned} g_{r,c} &= \alpha_{r,c} \sum_{m',c'} \varphi_{c,c'} \mathbf{Z}_{r,c',j^*} \\ \|g\|_2^2 &= \sum_{r,c} g_{r,c}^2 \\ \phi_m &= \sum_{r,c} \left( \sum_{m'} \left( K_{m,m'} \mathbf{X}^{(j)}_{m',r,c} \right) - \frac{1}{2}\lambda \right) g_{r,c}. \end{aligned}$$

In terms of these objects, it is straightforward to show that the relevant portion of the loss can be written as

$$\mathcal{L}(f) = \|g\|^2 \|Kf\|^2 - 2 f^T \phi + \cdots .$$

We would like to minimize this subject to the constraint that $f_m \geq 10^{-10}$. To approach this problem we use a projected gradient descent approach. We start by selecting a search direction, namely the gradient of the Lagrangian:

$$\Delta = \phi - \|g\|^2 K^T K f.$$

We then zero out the coordinates of this search directions which point negatively along the active constraints:

$$\tilde{\Delta}_m = \begin{cases} 0 & \text{if } f_m = 10^{-10} \text{ and } \Delta_m \leq 0 \\ \Delta_m & \text{otherwise.} \end{cases}$$

We then update $f$ by moving it somewhat in this search direction and then forcing it to be positive. How far should we move in the search direction? Following [13], we use the following carefully-chosen step-size:

$$f \leftarrow \left[ f + \frac{\phi^T \tilde{\Delta} - \|g\|^2 K^T K \tilde{\Delta}}{\left\| K \tilde{\Delta} \right\|^2} \tilde{\Delta} \right]_+ .$$

If we did not force $f$ to be positive, such updates would yield the best possible distance to travel along the search direction. However, due to the positivity-enforcement, one can find pathological examples where applying this update actually makes the loss worse. To be safe, we use a backtracking procedure; as long as the loss is made actively worse by this step, we cut the learning rate in half and try again.

$\varphi$ **update** Fix $c^*$. Let us look at the loss with respect to $\varphi_{c^*,1}, \varphi_{c^*,2} \cdots \varphi_{c^*,C}$. We find that it is given by

$$\mathcal{L}(\varphi_{c^*}) = \sum_{m,r} \left( \texttt{xmab}_{m,r,c^*} - \sum_{c'} \varphi_{c^*,c'} \left( \sum_{j,m'} \alpha_{r,c^*} \mathbf{K}_{m,m'} \mathbf{F}_{m',j} \mathbf{Z}_{r,c',j} \right) \right)^2$$
$$+ \lambda \sum_{c} \varphi_{c^*,c'} \left( \sum_{m,j} \alpha_{r,c} \mathbf{F}_{m,j} \mathbf{Z}_{r,c',j} \right) + \cdots .$$

Define

$$\Gamma_{c_1,c_2} = \sum_{m,r} \left( \sum_{j,m'} \alpha_{r,c^*} \mathbf{K}_{m,m'} \mathbf{F}_{m',j} \mathbf{Z}_{r,c_1,j} \right) \left( \sum_{j,m'} \alpha_{r,c^*} \mathbf{K}_{m,m'} \mathbf{F}_{m',j} \mathbf{Z}_{r,c_2,j} \right)$$
$$\phi_c = \sum_{m,r} \texttt{xmab}_{m,r,c^*} \left( \sum_{j,m'} \alpha_{r,c^*} \mathbf{K}_{m,m'} \mathbf{F}_{m',j} \mathbf{Z}_{r,c,j} \right) - \frac{1}{2} \lambda \left( \sum_{m,j} \alpha_{r,c*} \mathbf{F}_{m,j} \mathbf{Z}_{r,c,j} \right) .$$

Fixing all other variables, the problem of minimizing the loss with respect to $\varphi_{c^*}$ can then be understood as a quadratic programming problem.

$$\begin{aligned} \min_{\varphi_{c^*}} \quad & \frac{1}{2} \varphi_{c^*}^T \Gamma \varphi_{c^*} - \varphi_{c^*}^T \phi \\ \text{subject to} \quad & \varphi_{c^*} \geq 10^{-10} \end{aligned}$$

This problem is low dimensional and easy to solve using an off-the-shelf package. We use `scipy.optimize.nnls`.

#### L.1 Selecting $\omega$

So far, we have assumed that $\omega$ used in Equation 1 is a user-provided parameter. This $\omega$ represents the maximum tolerated reconstruction error. There are three methods for choosing this parameter which we can suggest:

**Interactively.** If the observation model is correct, the predicted values of $\mathbf{X}$ should be 'close' to the observed values. To discern this, our package provides an interactive method for selecting an $\omega$ which is satisfactory. This function starts with very large error tolerance (specifically we take $\omega$ to be half the maximum observed intensity. The function then allows the user to visually compare the true observations with the predicted values estimated with this value of $\omega$. If the predicted values appear to miss important features of the observed data, the user can then reduce $\omega$. The optimization will be re-run (warm-starting from the old initial condition, so this does not require very much time), and new predicted values are displayed. The function allows the user to continually reduce $\omega$ until the user deems that all the important features of the observed data are captured by the predicted values.

**Automatically.** An automatic method can be achieved by starting with the original data, slightly blur it, and take the average squared magnitude of the difference. This magnitude can be used to estimate the amount of speckle noise in the image. We can then choose $\omega$ so that the average reconstruction loss at each voxel is less than twice the value of this speckle noise.

**Via manually-labeled data.** If the user is willing to annotate a portion of their data with their beliefs about which rolonies are located at which positions, this annotated data can be used to select the $\omega$. Specifically, one can select the $\omega$ which yields the most accurate rolony detection.

In practice, we find that the interactive method is the most straightforward to use.
