## Supplemental Figures for "BARcode DEmixing through Non-negative Spatial Regression (BarDensr)"

S1 Fig

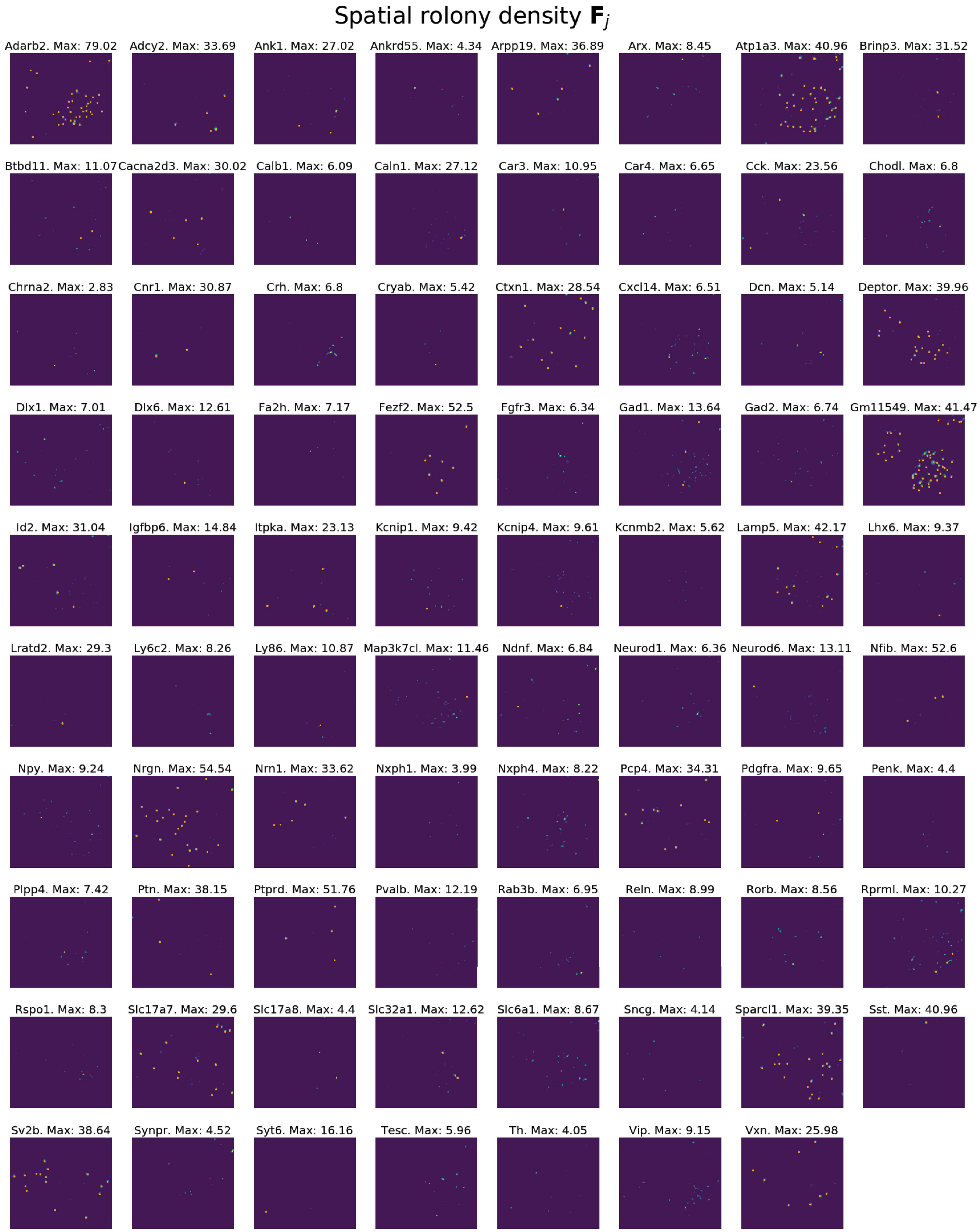
Spatial rolony density $\mathbf{F}_j$
Adarb2. Max: 79.02
Adcy2. Max: 33.69
Ank1. Max: 27.02
Ankrd55. Max: 4.34
Arpp19. Max: 36.89
Arx. Max: 8.45
Atp1a3. Max: 40.96
Brinp3. Max: 31.52
Btbd11. Max: 11.07
Cacna2d3. Max: 30.02
Calb1. Max: 6.09
Caln1. Max: 27.12
Car3. Max: 10.95
Car4. Max: 6.65
Cck. Max: 23.56
Chodl. Max: 6.8
Chrna2. Max: 2.83
Cnr1. Max: 30.87
Crh. Max: 6.8
Cryab. Max: 5.42
Ctxn1. Max: 28.54
Cxcl14. Max: 6.51
Dcn. Max: 5.14
Deptor. Max: 39.96
Dlx1. Max: 7.01
Dlx6. Max: 12.61
Fa2h. Max: 7.17
Fezf2. Max: 52.5
Fgfr3. Max: 6.34
Gad1. Max: 13.64
Gad2. Max: 6.74
Gm11549. Max: 41.47
Id2. Max: 31.04
Igfbp6. Max: 14.84
Itpka. Max: 23.13
Kcnip1. Max: 9.42
Kcnip4. Max: 9.61
Kcnmb2. Max: 5.62
Lamp5. Max: 42.17
Lhx6. Max: 9.37
Lratd2. Max: 29.3
Ly6c2. Max: 8.26
Ly86. Max: 10.87
Map3k7cl. Max: 11.46
Ndnf. Max: 6.84
Neurod1. Max: 6.36
Neurod6. Max: 13.11
Nfib. Max: 52.6
Npy. Max: 9.24
Nrgn. Max: 54.54
Nrn1. Max: 33.62
Nxph1. Max: 3.99
Nxph4. Max: 8.22
Pcp4. Max: 34.31
Pdgfra. Max: 9.65
Penk. Max: 4.4
Plpp4. Max: 7.42
Ptn. Max: 38.15
Ptprd. Max: 51.76
Pvalb. Max: 12.19
Rab3b. Max: 6.95
Reln. Max: 8.99
Rorb. Max: 8.56
Rprml. Max: 10.27
Rspo1. Max: 8.3
Slc17a7. Max: 29.6
Slc17a8. Max: 4.4
Slc32a1. Max: 12.62
Slc6a1. Max: 8.67
Sncg. Max: 4.14
Sparcl1. Max: 39.35
Sst. Max: 40.96
Sv2b. Max: 38.64
Synpr. Max: 4.52
Syt6. Max: 16.16
Tesc. Max: 5.96
Th. Max: 4.05
Vip. Max: 9.15
Vxn. Max: 25.98

S2 Fig

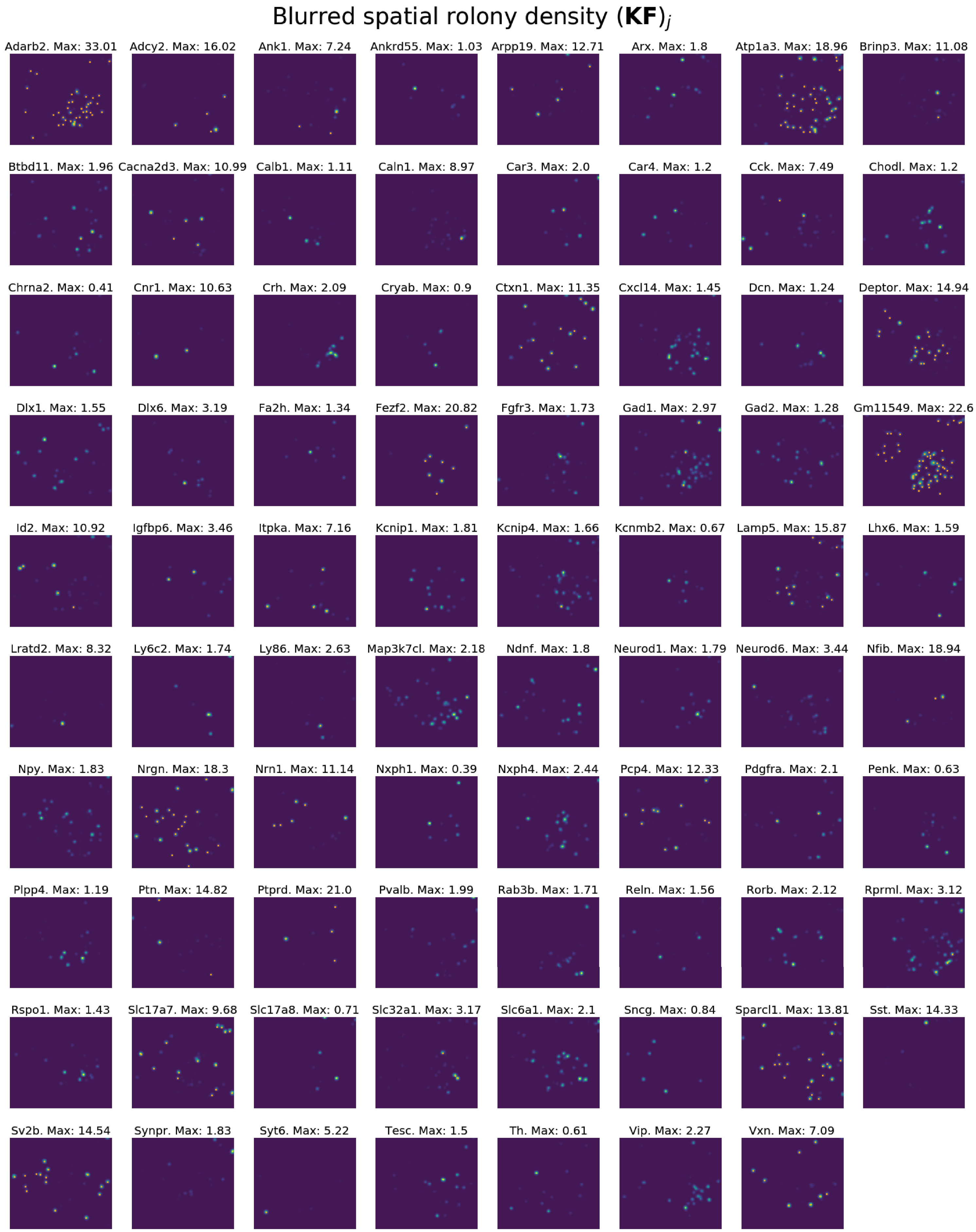

Blurred spatial rolony density $(\mathbf{KF})_j$
Adarb2. Max: 33.01
Adcy2. Max: 16.02
Ank1. Max: 7.24
Ankrd55. Max: 1.03
Arpp19. Max: 12.71
Arx. Max: 1.8
Atp1a3. Max: 18.96
Brinp3. Max: 11.08
Btbd11. Max: 1.96
Cacna2d3. Max: 10.99
Calb1. Max: 1.11
Caln1. Max: 8.97
Car3. Max: 2.0
Car4. Max: 1.2
Cck. Max: 7.49
Chodl. Max: 1.2
Chrna2. Max: 0.41
Cnr1. Max: 10.63
Crh. Max: 2.09
Cryab. Max: 0.9
Ctxn1. Max: 11.35
Cxcl14. Max: 1.45
Dcn. Max: 1.24
Deptor. Max: 14.94
Dlx1. Max: 1.55
Dlx6. Max: 3.19
Fa2h. Max: 1.34
Fezf2. Max: 20.82
Fgfr3. Max: 1.73
Gad1. Max: 2.97
Gad2. Max: 1.28
Gm11549. Max: 22.6
Id2. Max: 10.92
Igfbp6. Max: 3.46
Itpka. Max: 7.16
Kcnip1. Max: 1.81
Kcnip4. Max: 1.66
Kcnmb2. Max: 0.67
Lamp5. Max: 15.87
Lhx6. Max: 1.59
Lratd2. Max: 8.32
Ly6c2. Max: 1.74
Ly86. Max: 2.63
Map3k7cl. Max: 2.18
Ndnf. Max: 1.8
Neurod1. Max: 1.79
Neurod6. Max: 3.44
Nfib. Max: 18.94
Npy. Max: 1.83
Nrgn. Max: 18.3
Nrn1. Max: 11.14
Nxph1. Max: 0.39
Nxph4. Max: 2.44
Pcp4. Max: 12.33
Pdgfra. Max: 2.1
Penk. Max: 0.63
Plpp4. Max: 1.19
Ptn. Max: 14.82
Ptprd. Max: 21.0
Pvalb. Max: 1.99
Rab3b. Max: 1.71
Reln. Max: 1.56
Rorb. Max: 2.12
Rprml. Max: 3.12
Rspo1. Max: 1.43
Slc17a7. Max: 9.68
Slc17a8. Max: 0.71
Slc32a1. Max: 3.17
Slc6a1. Max: 2.1
Sncg. Max: 0.84
Sparcl1. Max: 13.81
Sst. Max: 14.33
Sv2b. Max: 14.54
Synpr. Max: 1.83
Syt6. Max: 5.22
Tesc. Max: 1.5
Th. Max: 0.61
Vip. Max: 2.27
Vxn. Max: 7.09

S3 Fig

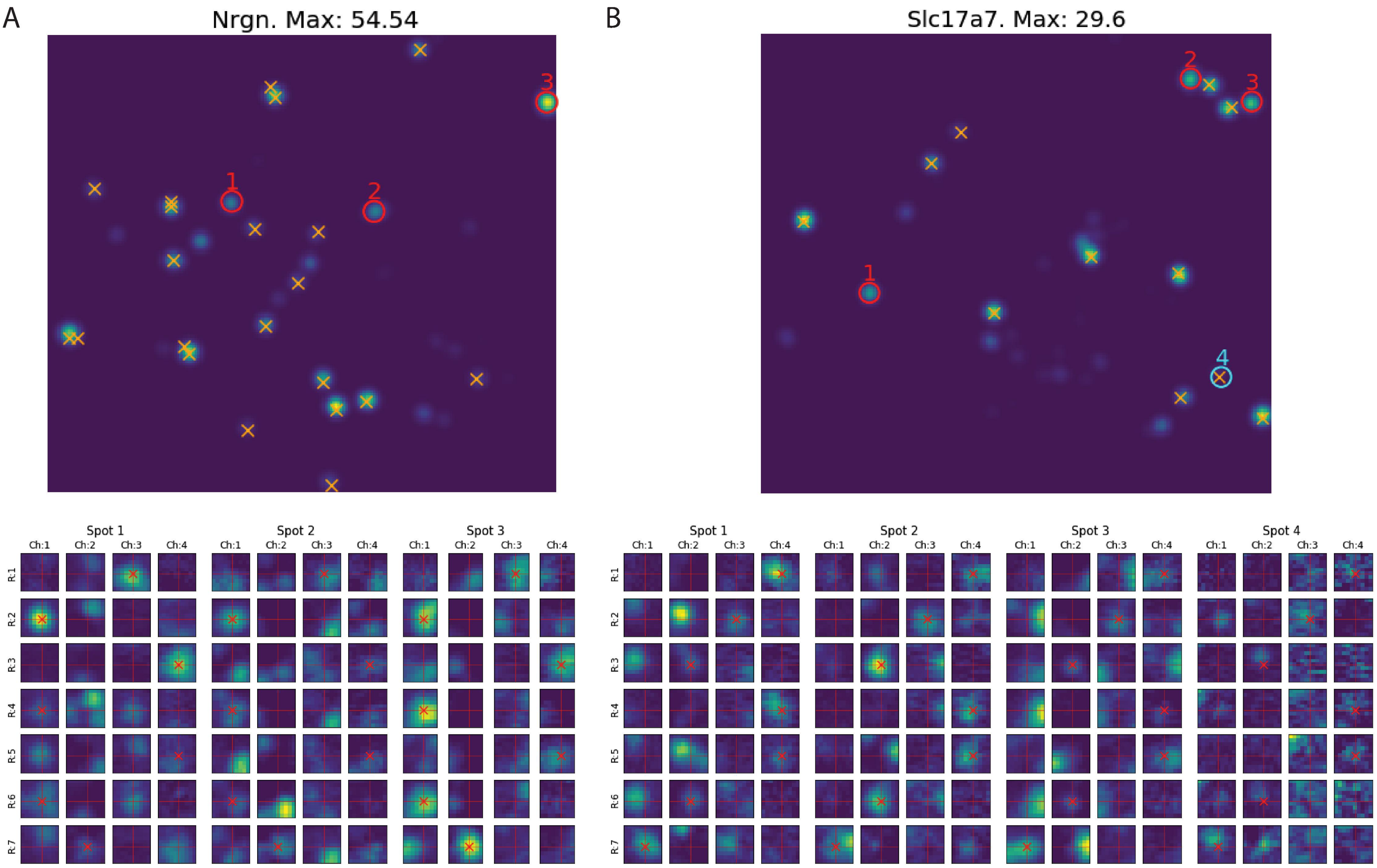
A
Nrgn. Max: 54.54
B
Slc17a7. Max: 29.6
Spot 1
Spot 2
Spot 3
Spot 4
Ch:1
Ch:2
Ch:3
Ch:4
R:1
R:2
R:3
R:4
R:5
R:6
R:7

S4 Fig

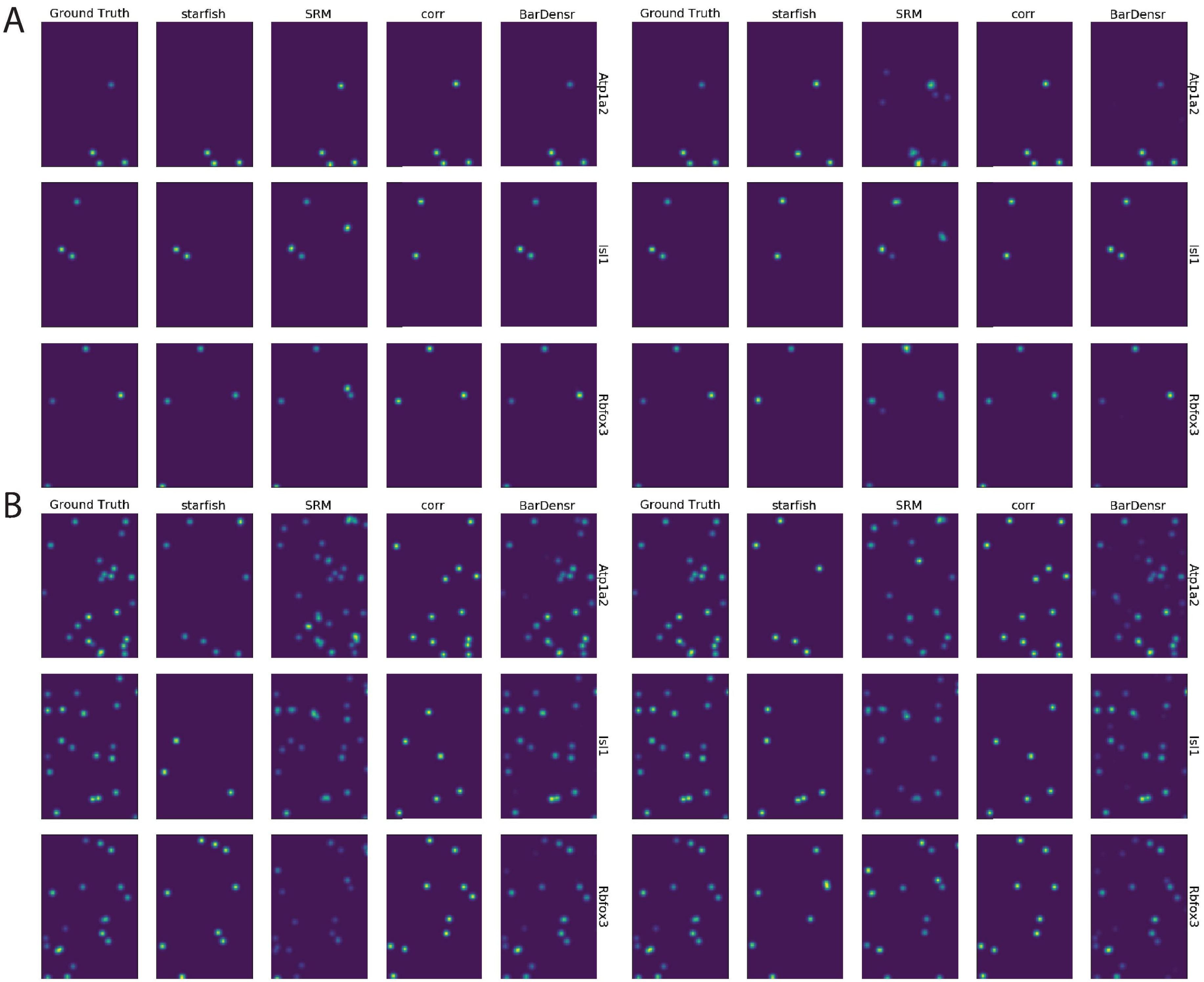
A
Ground Truth
starfish
SRM
corr
BarDensr
Atp1a2
Isl1
Rbfox3
B
Ground Truth
starfish
SRM
corr
BarDensr
Atp1a2
Isl1
Rbfox3

### S5 Fig

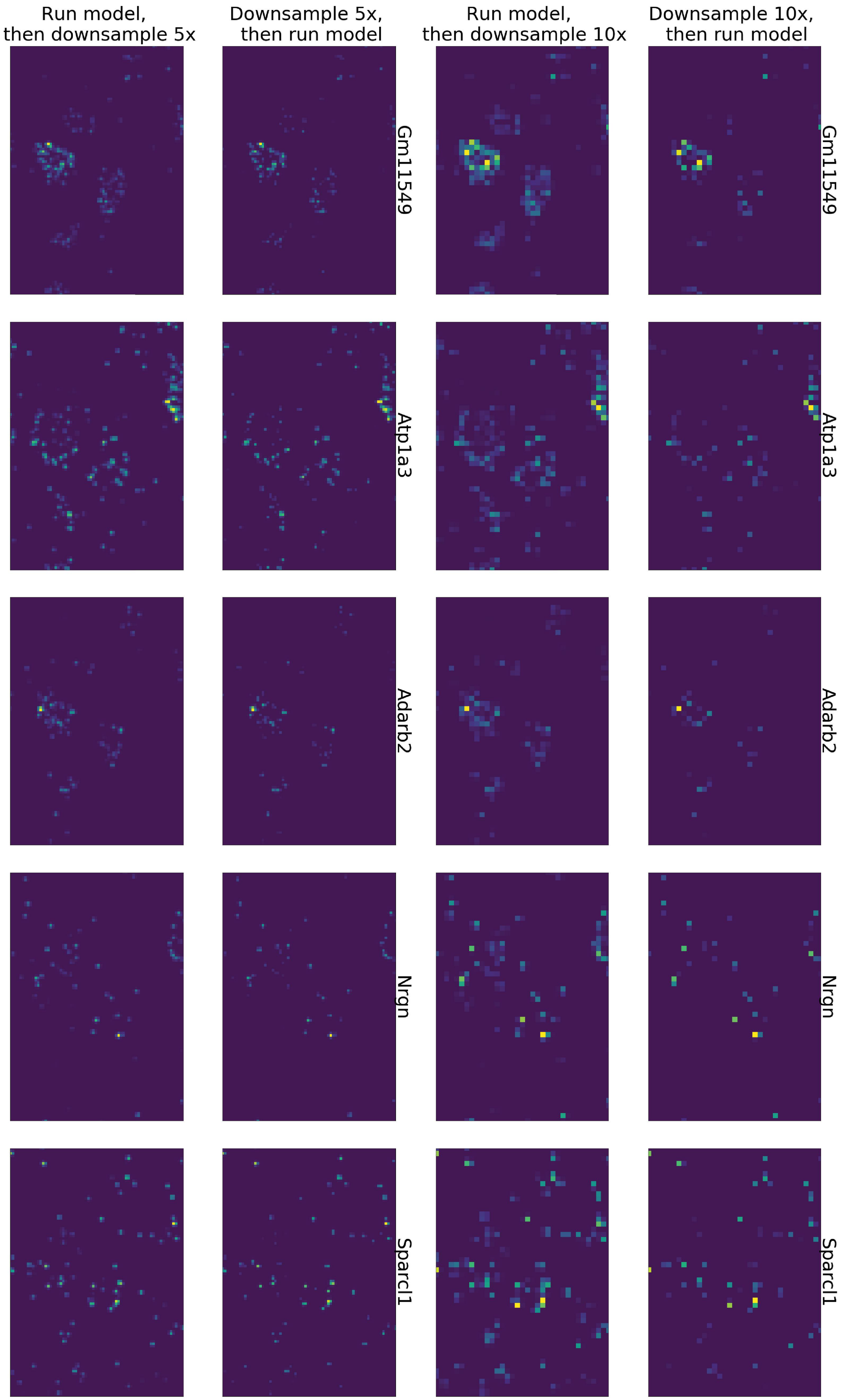
Run model, then downsample 5x
Downsample 5x, then run model
Run model, then downsample 10x
Downsample 10x, then run model
Gm11549
Atp1a3
Adarb2
Nrgn
Sparcl1
Gm11549
Atp1a3
Adarb2
Nrgn
Sparcl1

S6 Fig

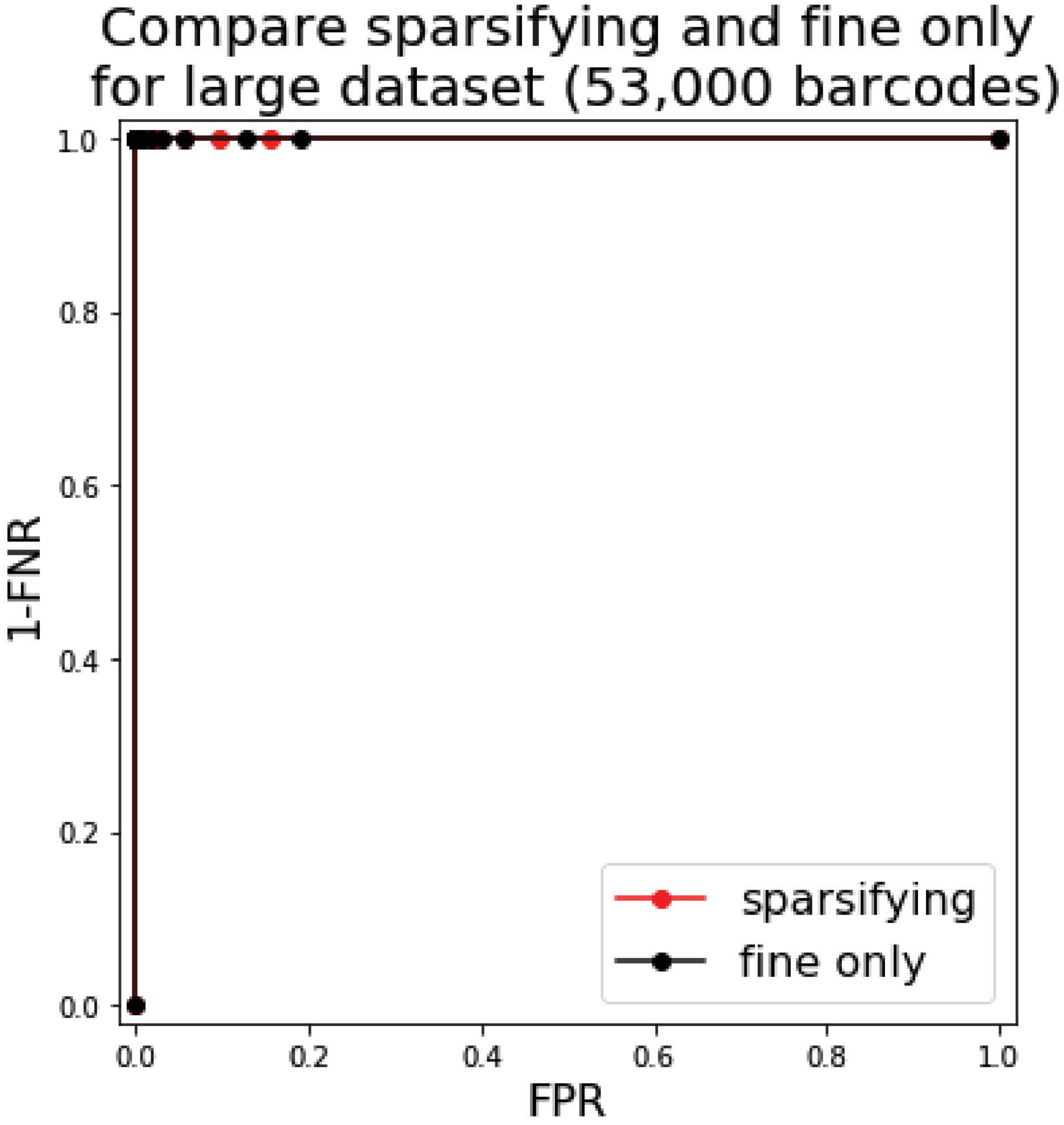
Compare sparsifying and fine only
for large dataset (53,000 barcodes)
1.0
0.8
0.6
0.4
0.2
0.0
1-FNR
0.0
0.2
0.4
0.6
0.8
1.0
FPR
sparsifying
fine only

S7 Fig

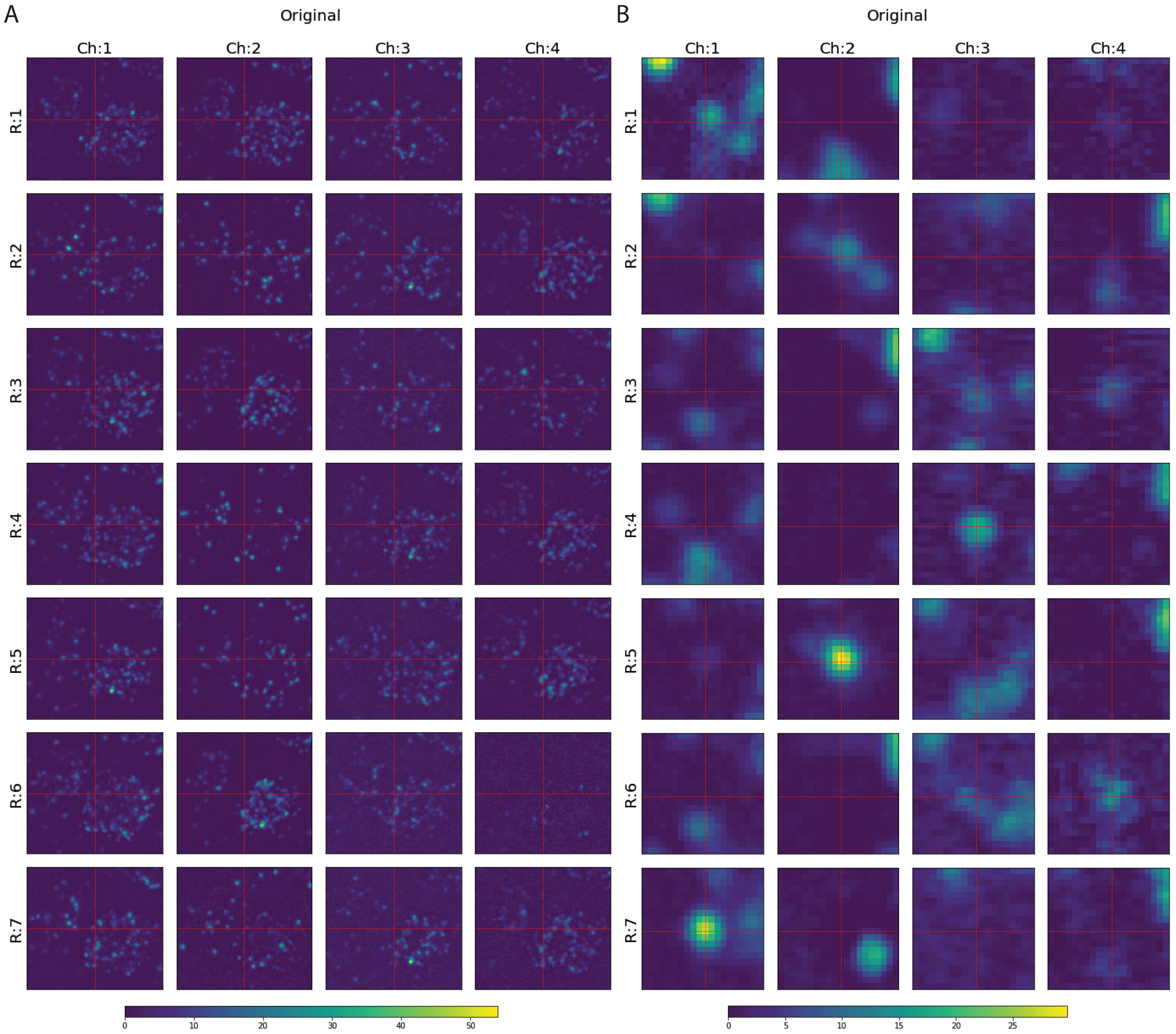
A
Original
Ch:1
Ch:2
Ch:3
Ch:4
R:1
R:2
R:3
R:4
R:5
R:6
R:7
0
10
20
30
40
50
B
Original
Ch:1
Ch:2
Ch:3
Ch:4
R:1
R:2
R:3
R:4
R:5
R:6
R:7
0
5
10
15
20
25

### S8 Fig

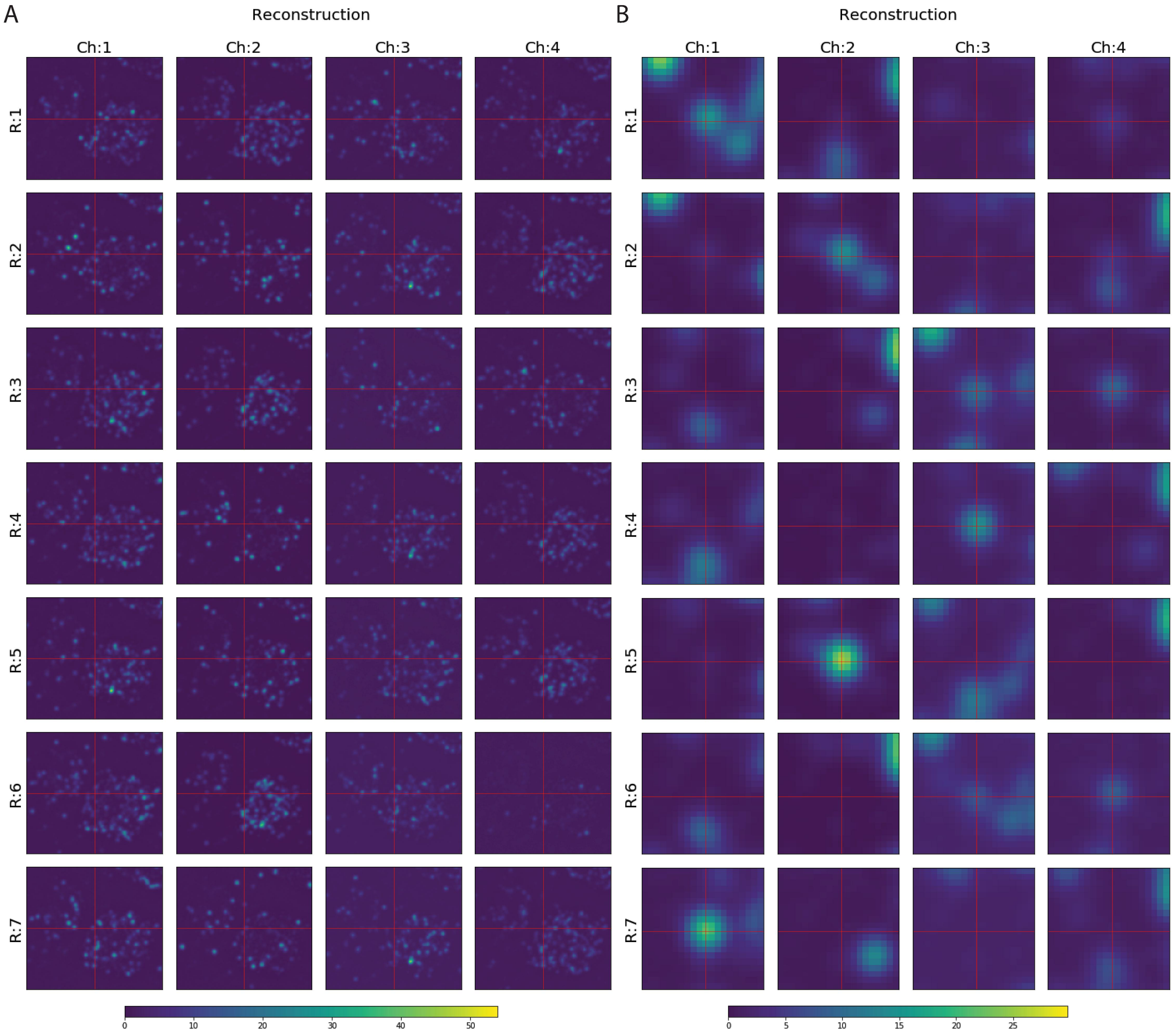

A
Reconstruction
B
Reconstruction
Ch:1
Ch:2
Ch:3
Ch:4
R:1
R:2
R:3
R:4
R:5
R:6
R:7
0
10
20
30
40
50
0
5
10
15
20
25

S9 Fig

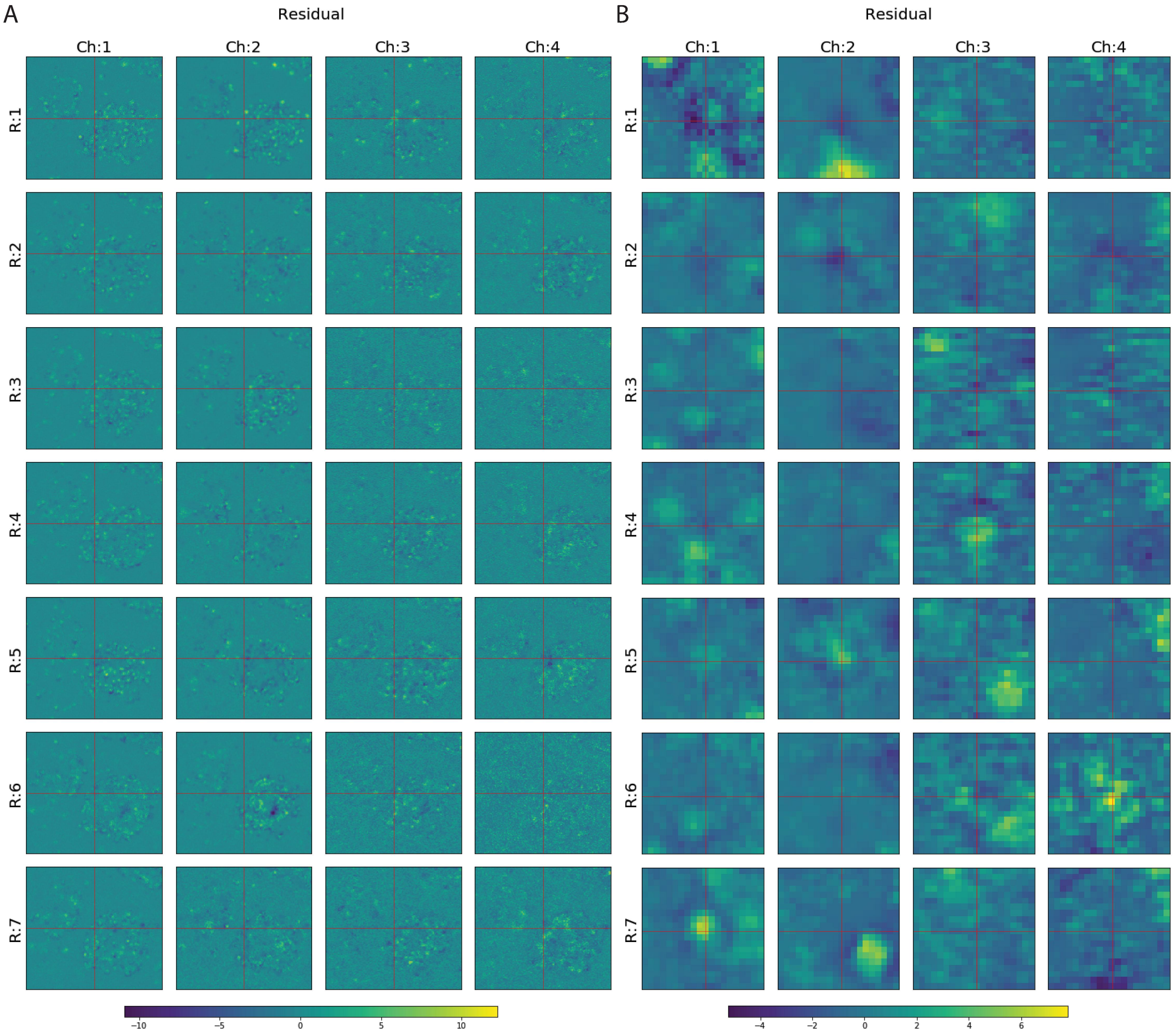
A
Residual
Ch:1
Ch:2
Ch:3
Ch:4
R:1
R:2
R:3
R:4
R:5
R:6
R:7
−10
−5
0
5
10
B
Residual
Ch:1
Ch:2
Ch:3
Ch:4
R:1
R:2
R:3
R:4
R:5
R:6
R:7
−4
−2
0
2
4
6

### S10 Fig

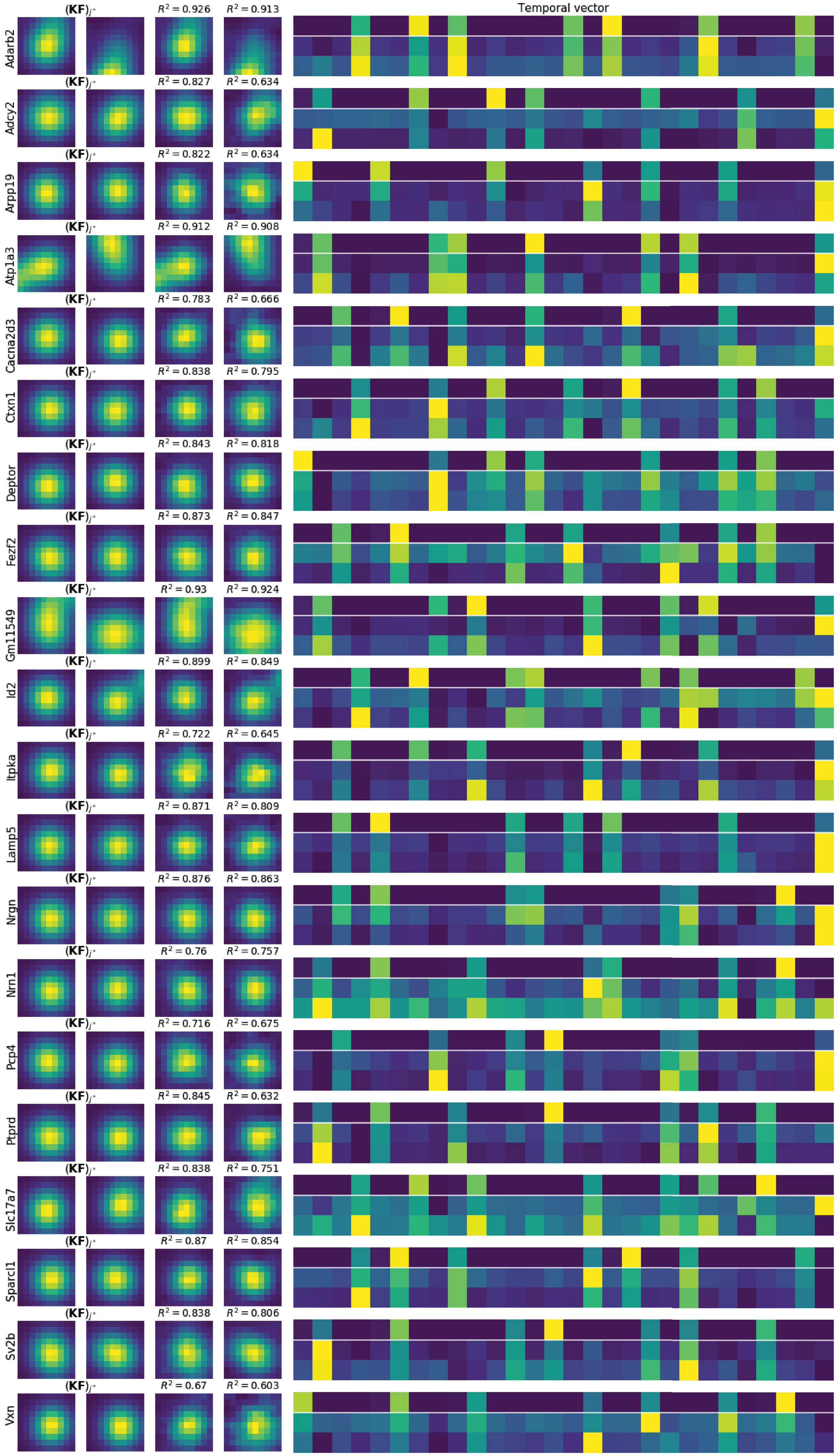
$(\mathbf{KF})_{j^*}$
Temporal vector
Adarb2
$R^2 = 0.926$
$R^2 = 0.913$
Adcy2
$R^2 = 0.827$
$R^2 = 0.634$
Arpp19
$R^2 = 0.822$
$R^2 = 0.634$
Atp1a3
$R^2 = 0.912$
$R^2 = 0.908$
Cacna2d3
$R^2 = 0.783$
$R^2 = 0.666$
Ctxn1
$R^2 = 0.838$
$R^2 = 0.795$
Deptor
$R^2 = 0.843$
$R^2 = 0.818$
Fezf2
$R^2 = 0.873$
$R^2 = 0.847$
Gm11549
$R^2 = 0.93$
$R^2 = 0.924$
Id2
$R^2 = 0.899$
$R^2 = 0.849$
Itpka
$R^2 = 0.722$
$R^2 = 0.645$
Lamp5
$R^2 = 0.871$
$R^2 = 0.809$
Nrgn
$R^2 = 0.876$
$R^2 = 0.863$
Nrn1
$R^2 = 0.76$
$R^2 = 0.757$
Pcp4
$R^2 = 0.716$
$R^2 = 0.675$
Ptprd
$R^2 = 0.845$
$R^2 = 0.632$
Slc17a7
$R^2 = 0.838$
$R^2 = 0.751$
Sparcl1
$R^2 = 0.87$
$R^2 = 0.854$
Sv2b
$R^2 = 0.838$
$R^2 = 0.806$
Vxn
$R^2 = 0.67$
$R^2 = 0.603$

S11 Fig

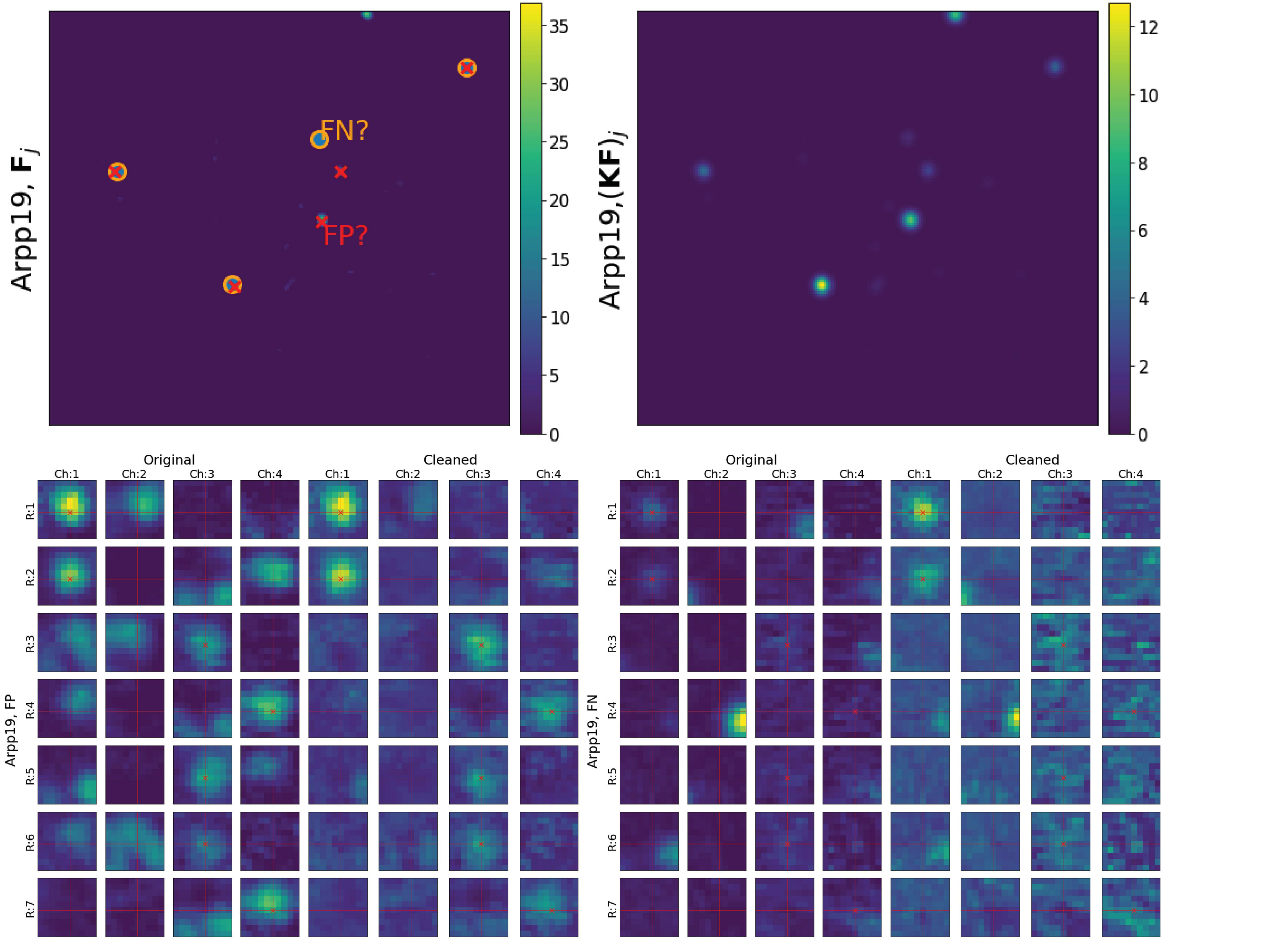

Arpp19, $\mathbf{F}_j$
FN?
FP?
35
30
25
20
15
10
5
0
Arpp19,$(\mathbf{KF})_j$
12
10
8
6
4
2
0
Original
Cleaned
Ch:1
Ch:2
Ch:3
Ch:4
R:1
R:2
R:3
R:4
R:5
R:6
R:7
Arpp19, FP
Arpp19, FN
